# Modulation of antigen delivery and lymph node activation in non-human primates by saponin adjuvant SMNP

**DOI:** 10.1101/2024.08.28.608716

**Authors:** Parisa Yousefpour, Yiming J. Zhang, Laura Maiorino, Mariane B. Melo, Mariluz A. Arainga Ramirez, Sidath C. Kumarapperuma, Peng Xiao, Murillo Silva, Na Li, Katarzyna K. Michaels, Erik Georgeson, Saman Eskandarzadeh, Michael Kubitz, Bettina Groschel, Kashif Qureshi, Jane Fontenot, Lars Hangartner, Rebecca Nedellec, J. Christopher Love, Dennis R. Burton, William R. Schief, Francois J. Villinger, Darrell J. Irvine

## Abstract

Saponin-based vaccine adjuvants are potent in preclinical animal models and humans, but their mechanisms of action remain poorly understood. Here, using a stabilized HIV envelope trimer immunogen, we carried out studies in non-human primates (NHPs) comparing the most common clinical adjuvant alum with Saponin/MPLA Nanoparticles (SMNP), a novel ISCOMs-like adjuvant. SMNP elicited substantially stronger humoral immune responses than alum, including 7-fold higher peak antigen-specific germinal center B cell responses, 18-fold higher autologous neutralizing antibody titers, and higher levels of antigen-specific plasma and memory B cells. PET-CT imaging in live NHPs showed that, unlike alum, SMNP promoted rapid antigen accumulation in both proximal and distal lymph nodes (LNs). SMNP also induced strong type I interferon transcriptional signatures, expansion of innate immune cells, and increased antigen presenting cell activation in LNs. These findings indicate that SMNP promotes multiple facets of the early immune response relevant for enhanced immunity to vaccination.

## INTRODUCTION

Adjuvants are essential for shaping the immune responses to subunit vaccines.^1,2^ The development of vaccines capable of neutralizing highly mutable pathogens may require adjuvants that can robustly promote germinal center GC responses where B cell selection and affinity maturation occur.^3–6^ To date, only a small number of adjuvants have been licensed for prophylactic vaccines in humans, and there is great interest in developing new vaccine adjuvants that can enhance GC responses, promote greater breadth of humoral immunity, or elicit broadly neutralizing antibody (bnAb) responses of greater magnitude and durability. Effective adjuvants may be particularly critical for promoting the development of bnAbs against HIV, influenza, sarbecoviruses and other emerging pathogens.

One particularly potent class of adjuvants are saponins, triterpene glycosides first studied as natural products isolated from the bark of the *Quillaja saponaria* tree. Saponins have surfactant-like activity and are typically formulated with lipids and cholesterol to avoid toxicity associated with membrane disruption caused by free saponin.^7^ Examples of licensed vaccines employing saponin adjuvants include Shingrix® and Mosquirix®, which employ Glaxo Smith Kline’s AS01 adjuvant^3^ (a liposomal formulation of saponin and the Toll like receptor (TLR)-4 agonist monophosphoryl lipid A (MPLA)), and the Novavax COVID-19 vaccine, which employs Matrix M adjuvant.^8^ Matrix M is an example of an Immune-Stimulating Complex (ISCOM) formulation of saponins, wherein saponins self-assemble with cholesterol and phospholipids to form ∼40 nm diameter cage-like nanoparticles. The precise mechanisms of action underlying the potency of saponin adjuvants remain poorly understood, though preclinical studies have demonstrated that saponins activate inflammasomes, promote antigen cross-presentation by dendritic cells, and trigger production of inflammatory cytokines and chemokines.^9–15^

We recently described a novel saponin adjuvant formed by physically incorporating the lipid TLR4 agonist MPLA into ISCOMs, forming saponin/MPLA nanoparticles (SMNP).^9,16^ SMNP was found to prime extremely strong germinal center (GC), T follicular helper (Tfh), and antibody responses in mice.^17–19^ Mechanistically, we found that in mice, SMNP enhanced lymph flow and the entry of antigens into draining lymph nodes (dLNs), accompanied by robust induction of inflammatory cytokines and chemokines in dLNs.^9^ In a pilot study employing a stabilized HIV envelope (Env) trimer immunogen in rhesus macaques, SMNP elicited stronger antibody titers, antibody effector functions, and tier 2 neutralizing antibody responses than two other experimental adjuvants of interest, a lipid-conjugated CpG TLR-9 agonist and a STING agonist.^9^ These preclinical experiments employed SMNP prepared using Quil A saponin, but based on the promising preclinical data, SMNP has been manufactured using highly purified QS-21 saponin for an ongoing first-in-humans clinical trial (HVTN144). However, it has remained unclear how QS-21-based SMNP would compare to the clinical gold standard adjuvant alum, and whether the mechanisms of action identified in rodents would also be relevant in a large animal model closer to humans.

Here, we compared immune responses and mechanisms of action following immunization of rhesus macaques with a stabilized HIV Env trimer immunogen termed MD39,^17^ adjuvanted either by aluminum hydroxide (Alhydrogel, hereafter alum for simplicity) or QS-21 SMNP (hereafter SMNP for simplicity). We found that SMNP elicited substantially stronger GC and Tfh responses than alum, as well as increased autologous neutralizing antibody titers. To gain insight into how the choice of adjuvant influenced early events in draining lymph nodes, we carried out studies combining PET imaging, flow cytometry, and scRNA-seq to evaluate antigen trafficking, immune cell recruitment and phenotype, and transcriptional status of immune cells in dLNs. Mirroring findings in rodents, these experiments revealed that, unlike alum, SMNP promoted antigen delivery to both proximal and more distal lymph nodes, triggered a strong type I interferon response in dLNs, and elicited changes in innate immune cells in both proximal and more distal lymph nodes of non-human primates (NHPs).

## RESULTS

### SMNP elicits much stronger germinal center responses than alum

We first carried out an immunogenicity study in rhesus macaques comparing alum with SMNP adjuvant. SMNP particles were synthesized incorporating QS-21 saponin, which is the highly purified saponin fraction employed in clinical saponin adjuvant formulations, to mimic the SMNP composition entering clinical testing.^3,11^ These adjuvants were combined with a highly stabilized native-like SOSIP HIV Env trimer termed MD39, representative of “polishing” immunogens in development for HIV vaccine regimens targeting the development of bnAbs.^17^ Two groups of rhesus macaques were immunized subcutaneously in the left and right inner thighs with MD39 trimer formulated with either alum or SMNP, administered at weeks 0, 8, 16, and 24 (**Figure 1A**). We opted to test this multi-boost immunization scheme motivated in part because we expected SMNP would be more potent than alum, but we wanted to determine if humoral responses elicited by alum could “catch up” to SMNP by simply increasing the number of immunizations. We first analyzed the kinetics of the GC responses by fine needle aspirate (FNA) sampling of draining inguinal LNs over the first 20 weeks of the study (**Figure S1A-C**). Previous studies have shown that FNAs are well-tolerated and provide a representative sample of the cellular composition of the entire LN.^18^ Alum immunization showed limited if any detectable Tfh expansion following each of the first 3 immunizations (**Figure 1B-C**). By contrast, SMNP elicited a pronounced expansion of Tfh cells at 2 weeks post prime immunization (WPPI) to a level ∼2.7-fold greater than the alum group, which contracted by wk 6, followed by less prominent changes at the subsequent boosts (**Figure 1B-C**). Similar trends were observed for total GC B cells over time: GC cells expanded to a peak of ∼10% of all B cells 2 WPPI for SMNP (**Figure 1D**). Interestingly, alum-induced GCs only began expanding after the third immunization (**Figure 1D**). Antigen-binding GC B cells were also assessed by staining B cells with tetramers of MD39 labeled with two different dyes (**Figure 1E**), and here the differences in GC responses were particularly striking: Alum immunization elicited weak trimer-binding GC responses that never exceeded 5% of the total GC B cell response, while SMNP elicited a trimer-specific GC B population that peaked at ∼11% of all GC B cells post prime, and further expanded to ∼25% of all GC B cells following the first boost at wk 8 (**Figure 1F**). Gating on GC B cells that stained brightly with antigen as a proxy for high antigen affinity (Antigen^high^ cells, **Figure 1E**), SMNP elicited a peak response where ∼3% of the trimer-specific cells were in this high-affinity gate (**Figure 1G**). Both the total GC response and trimer-specific response failed to expand as greatly following the third immunization, which we suspect reflects the high titers of antibody already present in the animals at the time of the third injection. Altogether, these data indicated that SMNP is capable of eliciting strongly enhanced GC responses in comparison to alum.

**Figure 1.**
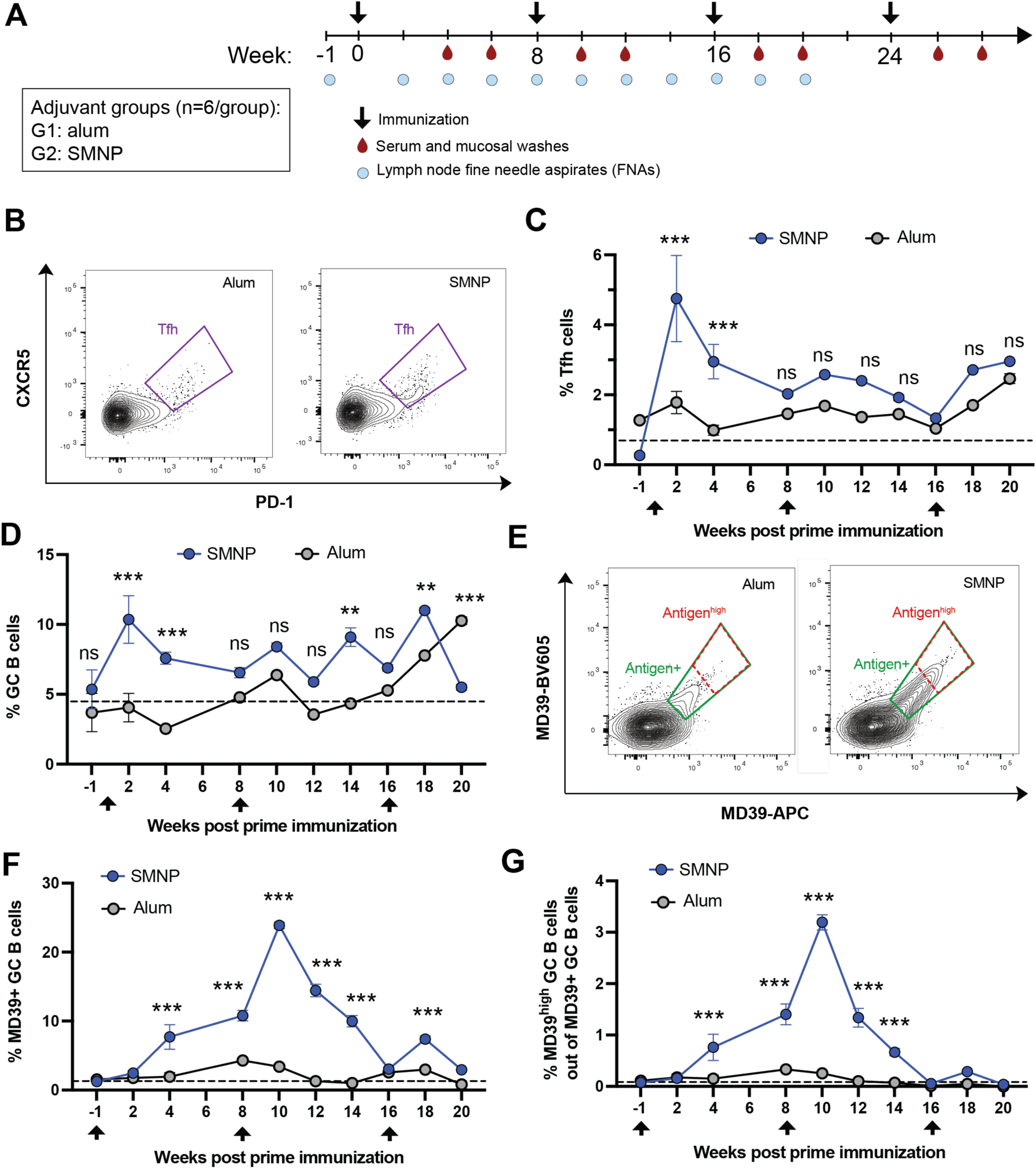
QS-21 SMNP promotes stronger Tfh and germinal center responses than alum. Rhesus macaques (*n* = 6/group) were vaccinated s.c. bilaterally in the inner thighs, with 50 µg MD39 trimer combined with either 500 µg alum or 188 µg SMNP per site at weeks 0, 8, 16, and 24. (A) Study diagram. (B-G) Tfh and GC responses detected by flow cytometry analysis of lymph node FNAs. (B) Representative Tfh flow cytometry plots gated on CD4^+^ T cells. (C, D) Frequencies of GC Tfh cells (CD3^+^CD4^+^PD1^+^ CXCR5^+^, C) and GC B cells (CD20^+^BCL6^+^Ki67^+^, D) over time. (E) Representative flow cytometry plots of trimer-binding GC B cells, indicating total antigen^+^ (green) and antigen^high^ (red) gates. (F-G) Frequencies of total (F) and high-binding (G) antigen-specific GC B cells. Dashed lines show baseline levels calculated from the responses at week −1. Statistical analyses were performed using one-way ANOVA, followed by Sidak’s multiple comparisons post-hoc test. (*P < 0.05, **P < 0.01, ***P < 0.001).

### SMNP primes higher antibody and bone marrow plasma cell responses than alum even following repeat dosing

We next evaluated early serum cytokine and plasmablast responses, serum antibody levels, and neutralizing antibody titers in the NHPs. Following the primary immunization and the first boost, we collected blood samples serially over 72 hr to monitor for elevations of systemic cytokines/chemokines. Alum immunization elicited very low levels of cytokines or chemokines post-prime or post-boost (**Figure S2**). By contrast, SMNP vaccination elicited transient spikes in IL-6, IL-10, CXCL10, and MCP1, which peaked by 24 hr and returned to baseline by 72 hr; IL-6 was particularly elevated following the boost (**Figure S2**). Transient inflammatory cytokine signatures in the blood are typical of strong adjuvants and are also reported in humans for saponin adjuvants.^19,20^

Antigen-specific plasmablasts in the peripheral blood were analyzed via a B cell ELISpot assay 6 days post each immunization. Weak antigen-specific IgG and IgA responses were detected following each immunization with alum, while SMNP induced robust priming of MD39-specific IgG and IgA plasmablasts following each booster immunization (**Figure 2A**). Both alum and SMNP immunizations primed high levels of MD39 trimer-binding serum IgG, but SMNP responses were higher, especially following the final boost at wk 24 (**Figure 2B**). Although IgA plasmablasts were detected early post-boosting with SMNP, serum IgA levels were very low for both immunizations (**Figure S3A**). Transient spikes in trimer-specific IgG were detected in nasal and vaginal washes post each boost that were not sustained, suggesting this was transudated antibody driven by peak antibody production post boost (**Figure S3B-G**). HIV autologous tier 2 neutralizing antibody titers were assessed after the first boost and after the final boost. Little neutralization was detected in either group after the first boost, but at wk 26 trimer immunization with SMNP had elicited mean neutralizing titers ∼ 1000, and these neutralizing titers remained ∼10-fold greater than that elicited by alum immunization at wk 28 (**Figure 2C-E**).

**Figure 2.**
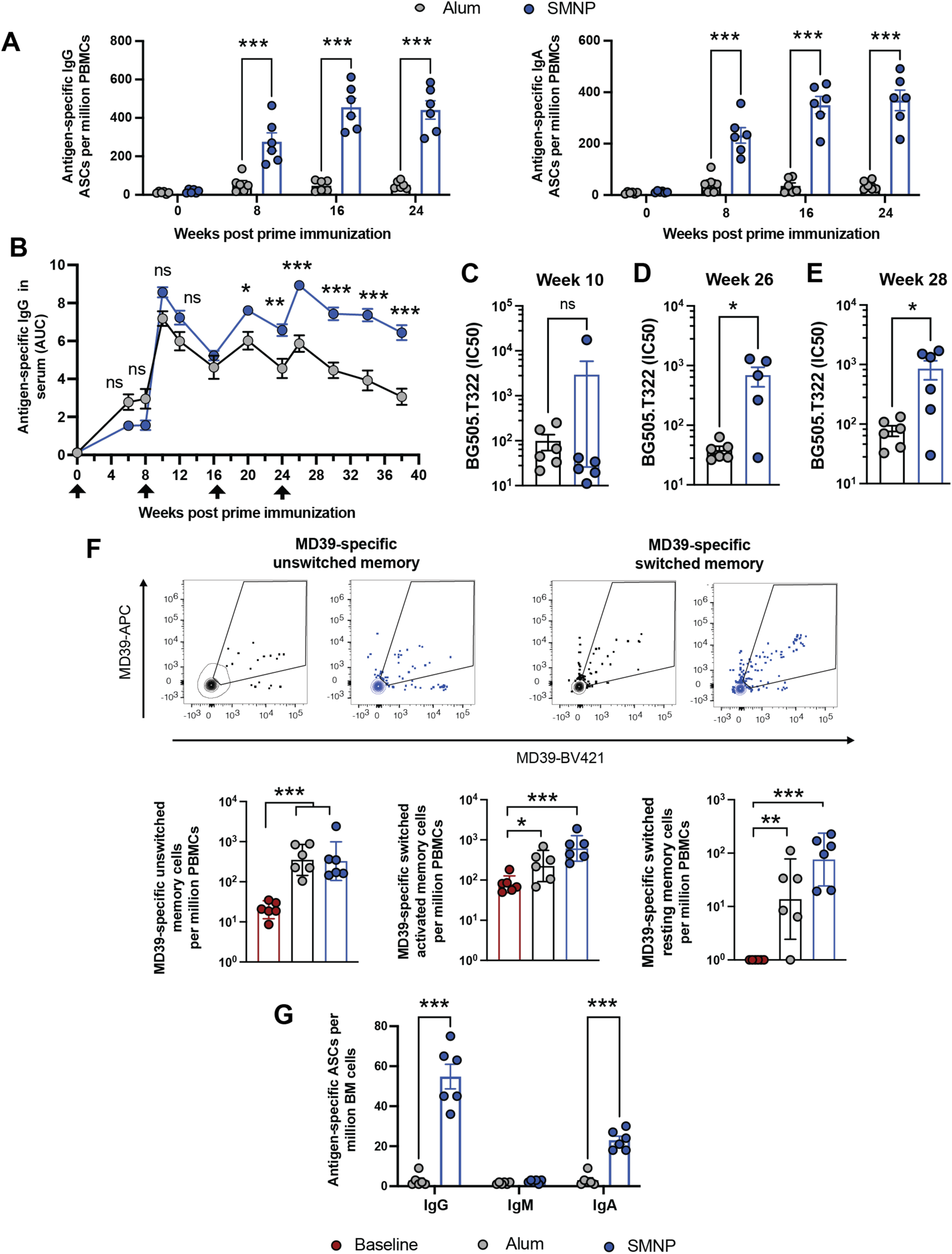
SMNP primes stronger plasmablast, serum antibody, and neutralizing antibody responses than alum. (A) Frequency of circulating antigen-specific antibody-secreting plasmablasts detected in PBMCs by ELISpot. (B) Serum IgG responses quantified as area under the curve (AUC) of ELISA optical density vs. serum dilution curves. (C-E) Autologous tier 2 neutralizing antibody titers at week 10 (C), 26 (D), and 28 (E) post prime immunization. (F) Antigen-specific memory B cells among PBMCs at week 34 post-vaccination. Shown are representative flow cytometry contour plots of antigen-specific cells gated on unswitched and switched memory B cells (top panels) and frequency of respective unswitched and switched (resting or activated) memory B cells (bottom panels). Data are presented as geometric mean ± 95% CI. (G) Trimer-specific IgM, IgG, or IgA antibody-secreting plasma cells were quantified at week 62 by ELISpot. Statistical comparisons were performed using one-way ANOVA, followed by Sidak’s (A, B) or Tukey’s post-hoc test (F) or Student’s *t* test (C, D, E, G). *, P < 0.05; **, P < 0.01; ***, P < 0.001. Log-distributed datasets (D, H) were log-transformed before statistical analysis.

Two key cellular outputs of the humoral response to vaccination are memory B cells and long-lived plasma cells. Flow cytometry analysis of peripheral blood mononuclear cells (PBMCs) at wk 34 to detect trimer-binding memory B cells revealed significant populations of class-switched resting and activated MD39-specific B cells (**Figure S4**),^21^ which trended higher for the SMNP-vaccinated animals (**Figure 2F**). Finally, we assessed plasma cell development by quantifying bone marrow (BM)-resident antibody secreting cells (ASCs) secreting different antibody isotypes by ELISPOT at wk 62. While alum elicited very low frequencies of trimer-specific ASCs, high levels of IgG- and clearly measurable levels of IgA-secreting plasma cells were detected in the SMNP group (**Figure 2G**). Thus, SMNP appears to be more effective than alum in promoting multiple facets of the humoral response to HIV Env immunogens.

### SMNP promotes antigen distribution to many lymph nodes in the draining lymphatic chain

Based on the findings from the immunogenicity study, we sought to gain insights into why SMNP was so much more effective in driving the humoral response than alum. In mice, we found that SMNP induced lymph vessel swelling, enhanced antigen drainage through lymph, and increased antigen entry into draining lymph nodes, leading to priming of GC responses in both proximal and secondary draining LNs.^9^ To determine if altered antigen trafficking could also underlie the efficacy of SMNP in NHPs, we carried out a study to visualize antigen dissemination from the injection site following alum or SMNP immunization using PET imaging in rhesus macaques. MD39 was conjugated with the radiometal chelator DOTA, to enable ^64^Cu labeling of the protein. Groups of macaques were then immunized s.c. in the left inner thigh with ^64^Cu-MD39 trimer mixed with alum or SMNP, and imaged serially over time by PET with tandem X-ray computed tomography (CT, **Figure 3A**). Co-registered PET/CT images revealed that after 24 hr, antigen was primarily localized at the injection site and the most proximal 2-3 draining inguinal and iliac lymph nodes for alum/MD39-immunized animals (**Figure 3B**, left). In striking contrast, by 24 hr, much of the antigen in the trimer/SMNP-immunized animals had left the injection site and accumulated in a series of 6-8 lymph nodes in the ipsilateral draining lymphatic chain (**Figure 3B**, right, and **Figure S5**). Quantitation of the mean Standard Uptake Values (SUVmean) at the site of injection (SOI) showed almost complete clearance of antigen in the SMNP group by 48 hr, while ∼50% of the antigen was still retained at the SOI in the alum group at this time point (**Figure 3C**).

**Figure 3.**
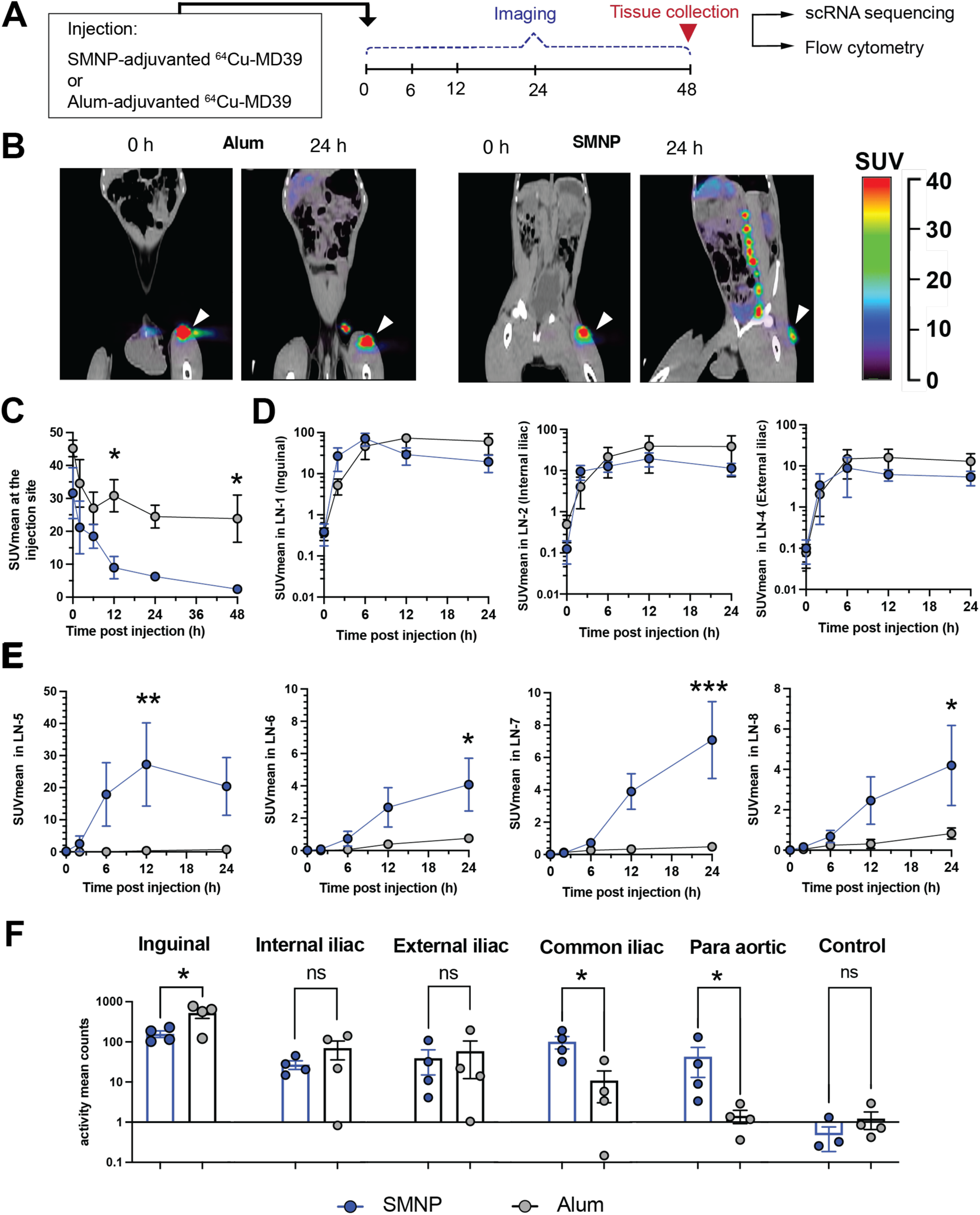
SMNP promotes antigen delivery to more distal lymph nodes than alum. NHPs (*n* = 4 animals/group) were immunized s.c. in the left inner thigh with 50 µg ^64^Cu-labeled MD39 trimer combined with either 500 µg alum or 188 µg SMNP, and then imaged over time by PET/CT scanning. (A) Timeline for PET/CT imaging studies. (B) Pseudocolor PET signal overlaid on grayscale CT scans at 0 and 24 hr post immunization. White arrows indicate injection site. (C-E) Radiolabeled MD39 signal from PET scan quantified as mean standardized uptake value (SUVmean) of gated ROIs at 0, 6, 12, 24, and 48 hr post injection. Shown are SUVmean of antigen signal over time at the injection site (C), most proximal draining lymph nodes (D), and more distal lymph nodes in the ipsilateral draining lymphatic chain, numbered in order from nearest to farthest from injection site (E). Data are presented as mean ± SEM. Statistical analyses were performed using one-way ANOVA, followed by Sidak’s post-hoc test. (*P < 0.05, **P < 0.01, ***P < 0.001). (F) SUVmean of PET antigen signal measured in the indicated lymph nodes *ex vivo* following tissue collection 48 hr post injection. Data are presented as mean ± SEM. Statistical comparisons were performed by Student’s t test (*P < 0.05).

Examining antigen uptake in the ipsilateral draining lymph node chain, we numbered the lymph nodes from 1 (nearest to the injection site, the inguinal LN) to 8 (most distal LN) and tracked the SUVmean over time for the nodes that could be definitively identified in animals over time. Antigen levels were highest in the 3 most proximal nodes (inguinal, internal iliac, and external iliac), and not statistically different for alum vs. SMNP (**Figure 3D**). However, in the more distal nodes, significant antigen uptake was still detected for trimers administered with SMNP, whereas negligible levels of signal were detected for trimers administered with alum (**Figure 3E**). At 48 hr the animals were sacrificed and LNs were collected for confirmation of signal by *ex vivo* PET scanning (**Figure 3A**); this analysis showed the same trend as the *in vivo* imaging, with the inguinal, internal iliac, and external iliac nodes acquiring similar levels of antigen for both alum and SMNP, while the next two more distal nodes (common iliac and paraortic) had 10-fold or more antigen signal in the SMNP group vs. the alum group (**Figure 3F**). (More distal nodes were too difficult to locate in the necropsy for further analysis). Altogether, these data suggest that, similar to observations in mice, SMNP promotes antigen trafficking to multiple lymph nodes, and antigen is seen reaching a series of nodes extending well beyond those most proximal to the injection site.

### SMNP induces a strong type I interferon signature in draining LNs

We next sought to assess changes in lymph node activation induced by SMNP vs. alum in the proximal inguinal draining lymph nodes. Lymph nodes from three NHPs per group were collected following PET imaging at 48 hours post immunization, and were processed for scRNA-seq analysis (**Figure 3A**). A contralateral inguinal lymph node without antigen signal from an NHP immunized with alum was collected as the control. Single cells, which were CD3^low^CD8^low^CD20^low^, were flow-sorted to moderately enrich for cells of myeloid lineage using flow cytometry (**Figure S6A**). After quality control of the scRNA-seq data, we identified seven cell lineages including natural killer (NK) cells, monocytes, macrophages, dendritic cells (DCs), T cells, B cells, and CD34^+^ hematopoietic progenitor cells, and recovered 8,785, 22,422, and 22,401 cells from the LNs of the control, alum, and SMNP groups, respectively (**Figure 4A-B**, **Figure S6B**).

**Figure 4.**
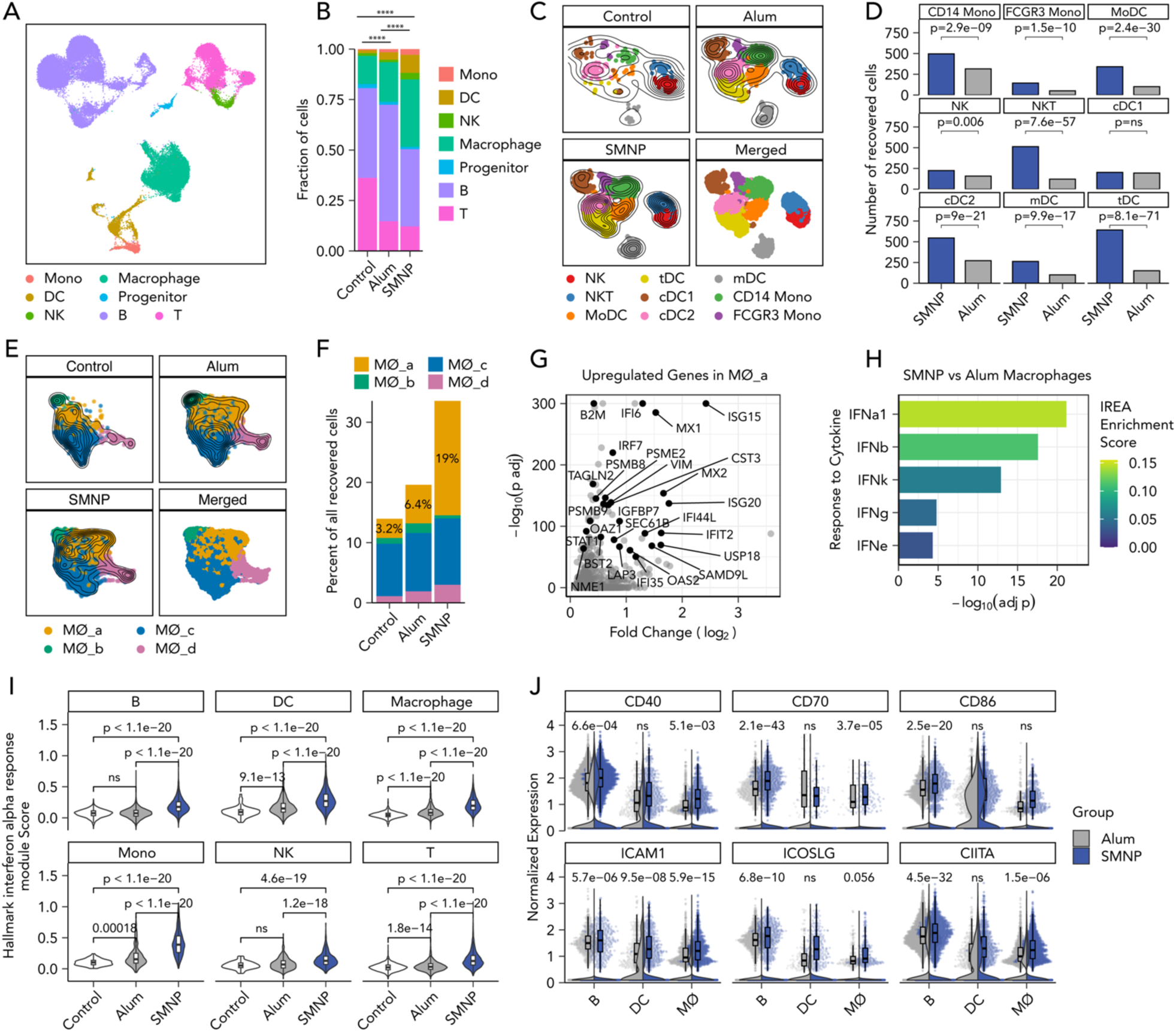
SMNP induces a strong type I interferon signature in draining lymph nodes. (A) UMAP of recovered cell lineages. (B) Distribution of recovered cell lineages in control contralateral (n_cell_=8785, n_sample_=1), alum (n_cell_=22422, n_sample_=3), and SMNP (n_cell_=22401, n_sample_=3) draining lymph nodes. P-values were calculated using a two-tailed chi-squared test with Bonferroni post-hoc correction. ****P < 0.0001 (C) UMAP of cell phenotypes among monocytes, dendritic cells (DCs), and NK cells. (D) Bar plots of the numbers of recovered cells between SMNP and alum for NK, DC, and monocyte phenotypes. P-values were calculated from Fisher’s exact test with Bonferroni post-hoc correction. (E) UMAP of phenotypic clusters of macrophages (MØs). (F) Percent of recovered macrophage clusters per group. For (A, C, and E), the contour lines represent the density of overlapping cells. (G) Volcano plot of significantly upregulated genes in the MØ_a cluster compared to the rest of the macrophage clusters. (H) Bar plot of significantly enriched cytokine responses in SMNP macrophages compared with alum macrophages based on the immune response enrichment analysis (IREA). (I) The cellular module scores for the Hallmark interferon alpha response gene set of individual cells from each group per cell lineage. P values were calculated using the Kruskal-Wallis test and Dunn’s post hoc test with Bonferroni post-hoc correction. (J) The expression levels of representative co-stimulatory genes in antigen-presenting cells (macrophage, DC, B cell) from SMNP (blue) and alum (grey) groups. The violin plots show the overall density of expression levels. The jittered dots represent individual cells and their expression levels. The boxplots show the distribution of cells with non-zero expression. P values were calculated using all cells with the Wilcoxon rank sum test with Bonferroni post-hoc correction.

For this very early time point, we focused on the analysis of innate cells. We identified two clusters of monocytes including *CD14*^high^ monocytes and CD16(*FCGR3*)^high^ monocytes, two clusters of NK cells, one of which is *CD3^high^KLRB1^low^* NKT-like, five clusters of DCs including migratory DCs (mDCs) with high expression of *CCR7*, *FSCN1*, *MARCKSL1*, and *LAMP3* ^22^, type 1 conventional DCs (cDC1), expressing *CLEC9A, CADM1, BATF3, XCR1, BTLA* ^23–25^, type 2 conventional DCs (cDC2) with canonical markers of *CD1C, FCER1A, CLEC10A*, and *CLEC4A* ^23–25^, monocyte-derived DCs (MoDC), expressing *NCOR, ZEB2*, and *SLC8A1* ^26–28^, and recently identified *AXL*^+^ transitional DCs (tDCs) ^29–33^, which are effective at priming CD4^+^ T cells, with a continuum of cDC2 and plasmacytoid DC-lineage markers such as *TCF4, CD141(THBD), SPIB, IRF7, BCL11A*, and *PPP1R14A* (**Figure 4C**, **Figure S6C**). Except for cDC1 cells, cell counts and frequencies of each innate cell population were significantly increased in SMNP LNs compared to alum LNs (**Figures 4D** and **S6D**). Particularly, 4.3, 3.4, and 4.3-fold more NK T cells, MoDCs, and tDCs were found in SMNP LNs (**Figure 4D**).

We next focused on macrophages, which increased in frequency moving from control to alum to SMNP lymph nodes (**Figure 4B**). Four phenotypic clusters were identified, and one of these, MØ_a cells, were present at 3-fold and ∼6-fold higher frequencies in the SMNP group vs. alum and control LNs, respectively (**Figure 4E-F**). Differential gene expression analysis of MØ_a cells vs. the other macrophage clusters showed that the former upregulated interferon (IFN)-responsive genes such as *ISG15, MX1, IFI6,* and *B2M*.^34^ (**Figure 4G**). This suggested that SMNP polarized macrophages more strongly toward a type 1 IFN-activated state than alum. To verify if this shift in phenotype was significantly different in SMNP vs. alum LNs, we performed a pseudo-bulk differential gene expression analysis between all macrophages from the SMNP vs. alum groups and identified 764 genes significantly upregulated (adjusted p-value < 0.01) in SMNP macrophages (**Figure S6E**). We next performed immune response enrichment analysis (IREA)^35^, where the upregulated genes in all macrophages from SMNP-vaccinated LNs were compared to gene signatures of macrophages polarized with 86 individual cytokines. We found that SMNP-treated macrophages were significantly enriched with gene signatures of response to IFN-α1, IFN-β, IFN-k, IFN-γ, and IFN3 cytokines (**Figure 4H**), indicating that SMNP indeed polarized macrophages towards an IFN-stimulated transcriptional state more strongly than alum.

To determine if type I IFN stimulation signatures were expressed in other cells isolated from SMNP-immunized lymph nodes, we scored the expression profile of each recovered cell by the Hallmark Interferon Alpha Response gene set from the Molecular Signatures Database (MSigDB)^36^. Strikingly, this analysis revealed that cells across all cell lineages upregulated genes involved in the type 1 IFN response (**Figure 4I**). To see if these gene expression signature extended to lymphocytes, we also analyzed the transcriptional phenotypes of B and T cells. We identified eight phenotypic clusters of B cells (**Figure S7A-B**) and five clusters of T cells (**Figure S7C-D**). Among the B and T cell clusters, we observed differences in the phenotypic composition between alum and SMNP and were able to identify clusters MBC_c and T_c of B and T cells, respectively, which were enriched in the SMNP group (**Figure S7E-F**). Both MBC_c and T_c cells showed strong type 1 IFN response signatures such as *ISG15, ISG20, MX1, IFI27, IRF9, B2M*, etc. (**Figure S7G-H**), corroborating that SMNP induced a broad IFN-stimulated state in LN cells.

Type 1 IFN signaling has been shown to enhance DC activation and differentiation^37^, improve T cell priming and the development of Tfh cells^38^, and promote B cell survival and differentiation into antibody-secreting cells^39,40^. Consistent with the literature, we found significantly increased expression of co-stimulatory molecules such as CD40, CD70, CD86, ICAM1, ICOSLG, and CIITA (the activator of MHC class II gene transcription) in antigen presenting cells from SMNP-vaccinated LNs compared to alum-vaccinated LNs (**Figure 4J**). Collectively, these results suggest that relative to alum, SMNP rapidly increased the frequency of diverse innate immune cells, triggered a broad type I IFN response, and promoted activation of antigen-presenting cells in draining lymph nodes.

### Innate immune activation induced by SMNP vs. alum in draining lymph nodes

To verify conclusions suggested by the scRNA-seq analysis, we carried out flow cytometry analysis on proximal inguinal and distal para-aortic lymph nodes recovered from NHPs post-PET imaging (**Figure S8**). Profiling of innate cells and B cells revealed that SMNP induced large increases in the number of neutrophils and CD68^+^ macrophages, and consistent trends toward increased numbers of CD123^+^ plasmacytoid and conventional DCs compared to alum in proximal draining inguinal LNs, changes that were not observed in control contralateral LNs (**Figure 5A** and **Figure S9A**). Similar patterns of expanded innate immune cell populations in SMNP-vaccinated animals were also measured in iliac and more distal para-aortic LNs (**Figure 5B**). We next examined markers of antigen-presenting cell (APC) activation. In proximal inguinal LNs, MHC II expression trended higher on DCs, monocytes, macrophages, and B cells in SMNP-vaccinated nodes vs. alum-vaccinated nodes, but did not reach statistical significance for any of these populations (**Figure 5C**). However, in more distal para-aortic LNs, MHC II was strongly upregulated on cDC2 cells and monocytes in response to SMNP but not alum (**Figure 5D-E**). We also examined CD80 expression on APCs, but no clear differences between groups were detected (**Figure S9B**). Finally, we carried out intracellular staining for expression of MX1, as an indicator of type I interferon signaling. We saw only a trend toward elevated MX1 expression in cDC2 and monocytes for in proximal inguinal nodes of SMNP-vaccinated animals (**Figure 5F**), but at the more distal para-aortic LNs, MX1 expression was significantly upregulated in these cell types (**Figure 5G-H**). These data suggest that SMNP not only increases antigen delivery to more distal lymph nodes, but also triggers innate immune stimulation necessary for promoting adaptive responses in these more distal sites.

**Figure 5.**
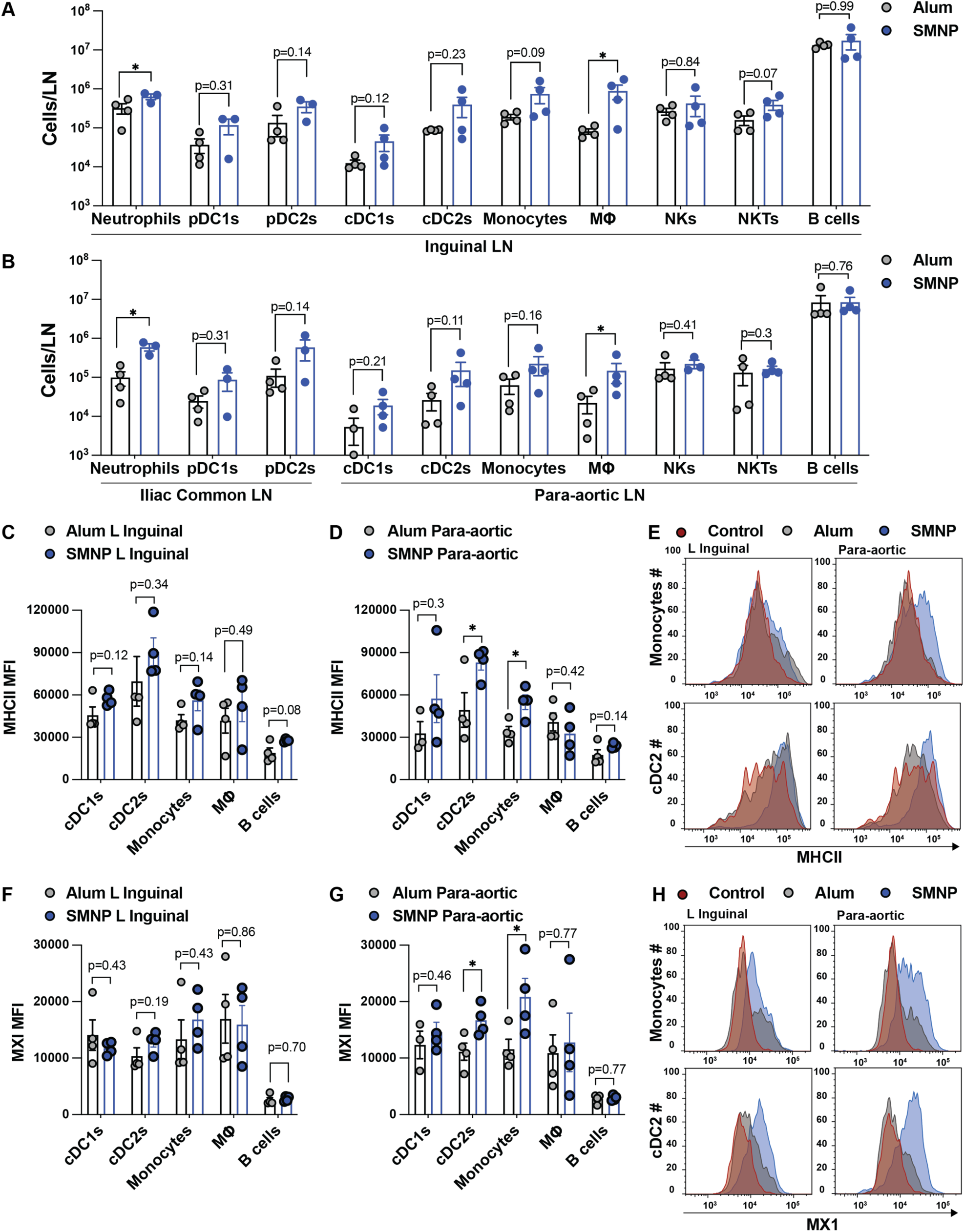
SMNP expands innate immune cell populations and triggers interferon signaling in both proximal and distal draining lymph nodes. LNs collected from NHPs following PET imaging at 48 hr post immunization with alum or SMNP adjuvant were processed for flow cytometry analysis. (A) Cell counts from proximal inguinal LNs. (B) Cell counts from iliac and distal para-aortic LNs. (C-E) MHC II expression on APC populations. Shown are MFI values for inguinal LN (C), para-aortic LN (D), and representative histograms of MHC expression on monocytes (E). (F-H) Intracellular straining for MX1 expression. Shown are MFI values for inguinal LN (F), para-aortic LN (G), and representative histograms of MX1 expression (H). *, P < 0.05 by Student’s *t* test.

## DISCUSSION

Adjuvants are critical components of vaccines that enhance protective immune responses, especially against poorly immunogenic antigens. Saponin adjuvants formulated with lipids have been shown to be safe for human use, and are used in both licensed vaccines and ongoing clinical trials.^41–44^ SMNP is a recently developed ISCOMs-type saponin adjuvant incorporating the TLR4 agonist MPLA, which has shown promising immunogenicity in diverse preclinical models,^45–49^ and recently entered first-in-humans testing in a phase I clinical trial (HVTN144, clinicaltrials.gov:NCT06033209). Here we performed a comparative analysis between SMNP and the traditional alum adjuvant in non-human primates, to evaluate the efficacy of this adjuvant in eliciting immune responses, and to further understand its mechanisms of action. SMNP for these studies was prepared using a highly purified saponin fraction termed QS-21, which is the form of saponin being used in the clinical trial of SMNP and is also the form of saponin employed in other saponin-based clinical adjuvant products such as GSK’s AS01b and ALFQ developed by the US Army.^41,42^ When evaluated with a native-like stabilized HIV Env trimer immunogen, we found that QS-21 SMNP elicited much stronger immunity than the gold standard alum adjuvant Alhydrogel, with greatly expanded GC and follicular helper T cell responses, elevated serum binding and neutralizing antibody titers, increased memory B cell production, and increased bone marrow plasma cell numbers. Mechanistically, we found through whole-animal and *ex vivo* PET/CT imaging that SMNP promoted antigen delivery to a greater pool of draining lymph nodes than alum, and induced a strong type I interferon signature associated with expanded innate immune cell populations and APC activation in both proximal and more distal dLNs.

One notable observation from the immunogenicity study was the significantly stronger antigen-specific GC response elicited by SMNP adjuvanted immunization, as determined by FNAs. GC B cells recognizing intact trimer expanded continuously over 8 weeks post priming immunization, and were further sharply escalated following the first boost. This was particularly noteworthy given the established relationship between early GC responses and the induction of neutralizing responses to HIV trimers.^50^ These early GC responses and output tier 2 neutralizing antibody titers primed by QS-21 SMNP were notably stronger than we observed in a parallel study comparing Quil A-based SMNP with alum and 3M-052 adjuvant,^45^ which might reflect increased potency of the purified QS-21 saponin used here. GC responses were more muted after the second boost, likely due to the high levels of pre-existing antibody titers by that time point.

The most striking observation made in these studies was the impact of SMNP adjuvant on the biodistribution of antigen following injection. In mice, we found that SMNP has multiple mechanisms of action that impact antigen trafficking, including induction of increased lymph flow, depletion of subcapsular sinus (SCS) macrophages, and augmented antigen entry in dLNs.^49^ These mechanisms correlated with induction of strong GC responses in both immediate draining and more distal dLNs in mice, but the small distances between LNs in rodents made it unclear if these effects would be observed in a large animal model closer to humans. Using PET/CT imaging to visualize antigen biodistribution in NHPs, we found that SMNP did indeed promote antigen dissemination to a greater number of LNs in macaques, with antigen reaching up to 8 LNs in the draining lymphatic chain. Importantly, we found evidence that SMNP was also activating innate immune responses across the LN chain, as innate immune cell populations were expanded and enhanced APC activation relative to alum was detected in both proximal and more distal LNs. We attempted to detect the induction of B cell priming across these dLNs, but at this very early timepoint of tissue sampling (48 hr), no clear antigen-specific B cell response could be measured.

NHPs are one of the closest-to-human animal models, especially in the pathophysiology of viral infections^51^. However, aside from the broad atlases of tissues from wildtype *Macaca fascicularis*^52^ and *Rhesus macaques*^53^, to our knowledge the impact of vaccine formulations on NHP LN myeloid cells at the single-cell transcriptomic level has not yet been examined. With scRNA-seq analysis, we found that even at the proximal LNs, SMNP induced a very distinct transcriptional program compared to alum. These findings echo transcriptional differences seen in an early microarray study profiling innate immune cells in the peripheral blood of NHPs vaccinated with 8 different adjuvants combined with HIV Env antigen^54^. Here the authors found that TLR4-containing formulations induced inflammatory and myeloid-associated transcriptional modules while ISCOMs led to IFN and antiviral modules^54^. Interestingly, SMNP, which is both an ISCOM saponin particle and carries a TLR4-agonist, induced a response that appears to be a hybrid of these two signatures– eliciting monocyte, DC, and NKT cells expansion in the LNs and strong type 1 IFN responses across all major cell lineages (**Figures 4D** and **4I**).

In summary, our studies demonstrate a strong immunogenicity of QS-21-based SMNP adjuvant in NHPs, with induction of novel mechanisms of action in NHPs that to our knowledge have not been described for other vaccine adjuvants. These results provide further insight into the potency of SMNP across preclinical animal models, and demonstrate some conservation of mechanisms of action between mice and larger animals. Importantly, QS-21 SMNP was also found to be safe in NHPs, eliciting minimal reactogenicity or injection site reactions, which will be a critical element for successful human use of this new adjuvant.

## MATERIALS AND METHODS

### Immunogen synthesis

The MD39 immunogen is a hyperstabilized BG505 SOSIP trimer and was synthesized as described previously.^55,56^ Briefly, trimer genes containing C-terminal His-tags were transfected into 293F cells grown in 293 Freestyle media (Life Technologies) by transient transfection with 293Fectin (Invitrogen). Protein was harvested from the culture supernatant at 5 days post transfection and purified by nickel affinity chromatography on a His-Trap HP column (Cytiva) followed by size exclusion chromatography using a HiLoad 16/600 Superdex 200 column (Cytiva). The purity and molecular weight of the purified trimer was confirmed by reducing and non-reducing SDS-PAGE.

### SMNP synthesis

SMNP is synthesized by self-assembly of dipalmitoylphosphatidyl choline (DPPC), monophosphoryl lipid A (MPLA), cholesterol, and saponin dissolved in MEGA-10 detergent at a 1:1:2:10 (DPPC:MPLA:Chol:Quil-A) mass ratio. Lipids, cholesterol, and synthetic MPLA (PHAD® Phosphorylated Hexaacyl Disaccharide) were purchased from Avanti Lipids and used as received. QS-21 saponin was obtained from Desert King and used as received. SMNP was synthesized as previously described,^49^ replacing Quil A saponin in the composition. The adjuvant was characterized by dynamic light scattering, HPLC, and a cholesterol quantification assay.

### Rhesus Macaques

Female Indian rhesus macaques (RMs, *Macaca mulatta*) were used in this study, aged between 3 and 4 years at time of first immunization. They were housed at the New Iberia Primate Research Center and maintained in accordance with NIH guidelines. The research protocols for these studies were reviewed and approved by the Institutional Animal Care and Use Committee (IACUC) of the University of Louisiana at Lafayette (Protocol #2017-8789-005).

### Macaque immunization study and sample collection

Groups of six female rhesus macaques were immunized subcutaneously in the caudal thigh muscle as at weeks 0, 8, 16, and 24 with 50 µg MD39 trimer formulated with 500 µg alum or 188 µg of SMNP, administered bilaterally in each thigh. For longitudinal titer studies, blood samples were collected by venipuncture from the femoral vein using serum collection tubes, frozen and stored at −80°C until analysis. Rectal and vaginal mucosal samples were collected using Merocel sponges as previously described^57^and stored at −80°C until analysis.

For PET imaging studies, MD39 trimer protein was labeled with DOTA-NHS-ester (Macrocyclics) in 0.1M sodium phosphate buffer pH 7.3 previously treated with Chelex 100 chelating resin (BioRad), and reacted at 4°C for 18 hr. Labeled trimer was purified using a Sephadex G-25 PD-10 Desalting column (GE) and stored at −80°C until use. ^64^Cu was obtained from the WIMR Cyclotron Labs at the University of Madison, Wisconsin, and DOTA-labeled MD39 trimer was complexed with the radiometal in chelexed 0.1 M sodium acetate buffer (pH 5.5) at 37°C with occasional mixing. Unbound Cu^64^ was removed from labeled MD39 by centrifugation through 10kDa microfilters (Amicon Ultra, UFC 5010BK) at 10,000x*g* for 10 min followed by 2 washes with pH 7.4 PBS, and the radioactivity measured in a dose calibrator. Two groups of 4 female rhesus macaques each were immunized subcutaneously in two cohorts with 50 µg ^64^Cu-labeled MD39 adjuvanted with either alum or SMNP. In the first cohort of animals, macaques were injected with 2.1 mCi ^64^Cu; in the second cohort, animals received 3.8 mCi ^64^Cu. These animals were then subjected to PET/CT scanning at 2, 4, 6, 12, 24 and 48 hours post vaccine administration.

### ELISA measurement of antibody titers

Anti-trimer titer was measured by ELISA. Nunc MaxiSorp plates were coated streptavidin overnight at 4°C and blocked with 2% BSA in PBS. Blocked plates were washed with washing buffer (PBS with 0.05% Tween-20) and incubated with Avi-tagged biotinylated antigen overnight at 4°C. Plates were washed again and serial dilutions of sera were added to plates and incubated the for 2 hr at room temperature, following which the plates were washed and incubated with anti-rhesus IgG- (and it subclasses), IgA-, and IgM-HRP (BioRad). ELISA plates were developed with TMB substrate and stopped by sulfuric acid 2 M. Absorbance values at 450 nm were measured, with a background correction wavelength of 540 nm. Serum titer was defined as the logarithm of the reciprocal of serum dilution that resulted in absorbance readings of more than 0.1 unit above background.

### Antigen-tetramer for trimer-specific B cell identification

To prepare MD39 probes, biotinylated MD39 trimer (Prepared by biotinylating Avi-tagged MD39 protein using the Avidity LLC BirA kit according to the manufacturer’s instructions) was mixed with either streptavidin-APC or streptavidin-BV605 in a 5:1 molar ratio. This mixture was then incubated for 20 minutes at room temperature to ensure proper tetramer formation before proceeding with the staining process.

### Flow cytometry analysis of germinal center responses

To longitudinally track the germinal center responses, lymph node fine needle aspirates (FNAs) were used to sample both right and left inguinal LNs identified to be the primary draining LNs by palpation. FNAs were conducted by a veterinarian. Samples were collected by inserting a 22-gauge needle attached to a 3mL syringe into the lymph nodes 4 times, and dispensed into the complete RPMI media (RPMI containing 10% fetal bovine serum, and 1ugml penicillin/streptomycin), and the needle was then flushed with media up to 3 times to collect all cells. Samples were centrifuged, and ACK (ammonium-chloride-potassium) lysis buffer was added if the sample had red blood cells. Cells were counted and cryopreserved in freezing medium (90% FBS, 10% DMSO) and stored in liquid nitrogen. For analysis, cells were thawed and dispersed through 70-µm cell strainers using the back of a 1-ml syringe plunger, and washed with PBS. The resulting single cell suspension was stained with Live/Dead Zombie Aqua for 15 min at room temperature, washed and then treated with antigen-tetramers for 30 min at room temperature. After pre-treatment with anti-CD16/32 Trustain FcX (10 minute at 4C) to block any nonspecific binding, cells were stained for 30 min at 4°C with antibodies (from BioLegend or BD Biosciences) against CD3, CD4, CD8, CD20, BCl-6, K67, PD1, and CXCR5. Antigen-specific staining was done using biotinylated trimer conjugated to streptavidin-BV421 (BioLegend) and streptavidin-APC (BioLegend). Antigen-specific GC B cells were gated as CD20+, CD3-, BCl-6+, Ki67+, MD39+ cells, and Tfh cells were gated as CD3+, CD20-, CD4+, CXCR5+, PD1+ cells.

### Flow cytometry analysis of memory B cell responses

Frozen PBMC samples were thawed and recovered in RPMI media with 10% FBS, supplemented with 1X penicillin/streptomycin. Fluorescent antigen probes were generated by mixing small incremental volumes of fluorophore-conjugated streptavidin (Streptavidin-BV421 and Streptavidin-APC) with biotinylated antigen in 1x PBS at 25°C over 45 min. Zombie UV fixable viability dye (Biolegend) was used according to manufacturer’s protocol. Cells were washed and incubated with Human TruStain FcX (Fc Receptor Blocking Solution) for 15 minutes at 25°C, followed by incubation with antigen probes for 30 min at 4 °C and then with surface antibodies for an additional 30 min at 4 °C. The staining antibody cocktail included: anti-human CD27 BUV395 (O323, BD Biosciences,1:20), CD20 BUV737 (L27, BD Biosciences, 1:20 dilution), IgM FITC (G20-127, BD, 1:20), IgD PE (Southern Biotech, 1:50), CD21 PE-Cy7 (B-ly4, BD, 1:20), CD14 APC-Cy7 (M5E2, Biolegend, 1:20), CD16 eFluor780 (3G8, Invitrogen, 1:20), CD3 APC-Cy7 (SP34-2, BD Biosciences, 1:20). After staining cells were fixed with 4% paraformaldehyde for 20 minutes at 4°C. Samples were spiked with Precision Count Beads and cell numbers were calculated according to the manufacturer’s protocol. At least 500,000 events per sample were acquired on an FACSymphony A3 (BD Biosciences) and data was analyzed using FlowJo v10 (FlowJo Inc).

### Bone marrow ELISPOT

Antibody-secreting cells (ASCs) in the bone marrow (BM) were assessed by enzyme-linked immune absorbent spot (ELISpot) assay. Briefly, 96-well multiscreen HTS filter plates (MilliporeSigma, catalog no. MSHAN4B50) were coated overnight at 4°C with 10 μg/ml of anti-monkey IgG, IgM or IgA (H&L) goat antibody (Rockland) or with 2 μg/ml of MD39 trimer for total or antigen-specific ASCs in BM, respectively. The plates were washed 4 times with PBS-0.05% Tween 20 (PBS-T) and 4 times with PBS and blocked with complete RPMI media for 2 h in a 5% CO2 incubator at 37°C. The BM cells were collected from the iliac crest of the animal in EDTA and separated by ficoll gradients method. Cells were counted and resuspended in complete RPMI media as 1 million per ml for total, and 5 million per ml for antigen-specific, plated in serial 3-fold dilutions, respectively, and incubated overnight in a 5% CO2 incubator at 37°C. Plates were then washed 4 times with PBS-T, and incubated with 1:1,000 diluted (PBS-T with 1% FBS) of anti-monkey IgG-, IgM-, or IgA-biotin-conjugated antibodies (Rockland), respectively, for 2 h at room temperature. The plates were again washed 4 times with PBS-T followed by addition of avidin D-horseradish peroxidase (HRP) (Vector Laboratories) diluted 1:1,000 in PBS-T with 1% FBS, for 1 h at room temperature. After washing 4 times with PBS-T and 4 times with PBS, plates were developed using the AEC substrate kit (BD Biosciences). To stop the reaction, plates were washed extensively with water followed by air drying. Spots were imaged and counted using the Immunospot ELISPOT Analyzer (Cellular Technology Limited). The number of spots specific for each Ig isotype was reported as the number of either total or antigen-specific ACSs per million BM cells.

### PET/CT imaging and Analysis

All whole-body PET/CT scans were performed on a Philips Gemini TF64 scanner. Acquisitions were done in 3D mode with an axial field of view of 57.6 cm. The PET and CT data were exported as DICOM files and analyzed using the Multi-Image Analysis GUI (MANGO, Research Imaging Institute, UT Health San Antonio). Decay-adjusted PET images (corrected to the time of vaccine injection) were normalized by the injected doses and body weight to generate Standardized Uptake Value (SUV) maps. To designate Regions of Interest (ROIs), spherical outlines were placed on target tissues, freehand outline drawings were made, and/or tissue contours on the SUV maps were identified through signal thresholding. These ROIs were then subjected to statistical evaluations to determine SUVmax, SUVmean, and SUVsum. The three-dimensional (3D) image data were exhibited as color-coded maximum intensity projections (MIPs) of the SUVmax maps. Analysis of the distal lymph nodes was performed by drawing 3D ROIs for the injection site and the most distal lymph node with a significant detectable signal (>10% of the maximum signal); these were subsequently projected onto a 3D surface-rendered PET image. The vaccine’s travel distance was measured by drawing a straight line on the 3D image between the SUVmax points of the injection site and the relevant lymph node.

Following the 48-h time point PET imaging session, the animals were euthanized in a humane manner with an intravenous administration of 200 mg/kg Beuthanasia-D®. An experienced veterinary pathologist then performed necropsies, and selected lymph nodes underwent a secondary PET imaging using the protocol described above.

### Flow cytometry for innate cells and B cells

Lymph nodes of vaccinated macaques were isolated 48 hours post-immunization, at the completion of PET imaging. The collected lymph nodes were first scanned before being mechanically dissociated into single cell suspensions. Cells were filtered, counted, and resuspended in freezing medium before storage in liquid nitrogen for subsequent analysis by flow cytometry and single cell RNA sequencing.

For flow cytometry analysis, Zombie UV fixable viability dye (Biolegend) was used according to manufacturer’s protocol. Cells were washed and incubated with Human TruStain FcX (Fc Receptor Blocking Solution) for 15 minutes at 25°C, followed by staining with a cocktail of fluorescent antibodies for 30 minutes at 4°C (panels 1-3 described below). After staining cells were either fixed with 4% paraformaldehyde for 20 minutes at 4°C (panel #2 and #3) or processed for intracellular staining with BD Cytofix/Cytoperm Fixation/Permeabilization Kit, according to the manufacturer’s instructions (panel #1). Intracellular staining was carried out in BD Perm/Wash buffer for 30 minutes at 4°C. After staining, samples were spiked with Precision Count Beads and cell numbers were calculated according to the manufacturer’s protocol. At least 500,000 events per sample were acquired on an FACSymphony A3 (BD Biosciences) and data was analyzed using FlowJo v10 (FlowJo Inc).

Panel #1 included anti-human CD16 BUV396 (3G8, BDBiosciences, 1:20), CD14 BUV737 (M5E2, BD Biosciences, 1:20), CLEC9A BV421 (3A4, BD Biosciences, 1:10), CD80 BV650 (L307.4, BD Biosciences, 1:20), HLA-DR PE (Tu36, Biolegend, 1:20), CD11c PE-Cy7 (3.9, Biolegend, 1:20), CD68 AF467 (eBioY1/82A, ebioscience, 1:10), CD20 APC-Cy7 (L27, BD Biosciences, 1:20). Intracellular staining: anti-human MX1 CoraLite® Plus 488 (Invitrogen, 1:20).

Panel #2 included anti-human CD16 BUV396 (3G8, BD Biosciences, 1:20), CD123 BV421 (6H6, Biolegend, 1:20), CD80 BV650 (L307.4, BD Biosciences, 1:20), CD14 AF488 (M5E2, Biolegend, 1:20), HLA-DR PE (Tu36, Biolegend, 1:20), CD11c PE-Cy7 (3.9, Biolegend, 1:20), CD66abce APC (TET2, Miltenyi, 1:20), and CD20 APC-Cy7 (L27, BD Biosciences, 1:20). Panel #3 included anti-human CD8 BUV396 (RPA-T8, BD Biosciences, 1:20), CD20 BUV737 (L27, BD Biosciences, 1:20), CD80 BV650 (L307.4, BD Biosciences, 1:20), NKG2A-FITC (REA110, Miltenyi, 1:20), CD14 PerCP/Cy5.5 (M5E2, Biolegend, 1:20), HLA-DR PE (Tu36, Biolegend, 1:20), CD3 APC-Cy7 (SP34-2, BD Biosciences, 1:20).

### Single-cell RNA sequencing preparation and analysis

Proximal inguinal LNs were collected from NHPs at 48 hr post immunization and processed into single cell suspensions as described above. Cells were stained for viability (Zombie NIR) and with antibodies against CD3 (APC-Cy7, SP34-2), CD8 (APC-Cy7, RPA-T8), CD20 (APC-Cy7, L27), HLA-DR (PE, Tu36). Live CD3^low^CD8^low^CD20^low^ cells were sorted on a BD FACS Aria (BD Biosciences) cell sorter, processed immediately using the commercial 5’ Single Cell GEX v2 platform (10x Genomics) following the manufacturer’s protocols, and sequenced on a NovaSeq 6000 system (Illumina).

Sequenced FASTQ files were aligned to the Rhesus macaques Mmul_10 genome assembly (Ensembl.org) using the cellranger cloud analysis pipeline (10x Genomics, v7.1.0) with default settings. Downstream analysis was performed in R (v4.3.2). The gene expression count matrices were processed and analyzed using Seurat (v5.0.1). ^58^ The initial quality control filtered out genes that were detected in less than 3 cells and removed cells with less than 500 genes, greater than 30000 UMI counts, and greater than 10% mitochondrial genes. Cells were log-normalized using the NormalizeData function, and variable genes were identified using the FindVariableFeatures function. The ScaleData function was used to scale the data to unit variance using a Poisson model and regress out RNA feature counts and percent of mitochondrial genes before performing principal component analysis (PCA) using the RunPCA function. Data integration was performed using the IntegrateLayers() function with the Harmony algorithm (v1.2.0)^59^. Thirty principal components (PCs) were used for constructing the nearest-neighbor graph with the FindNeighbors function. Thirty neighboring points and thirty PCs were used to generate uniform manifold approximation and projection (UMAP) with the RunUMAP function. Unsupervised clustering was determined using Louvain clustering as implemented in the FindClusters function. Differential gene expression analysis was performed using FindMarkers function. Clusters with strong expressions of marker genes associated with more than one cell population were denoted as doublet clusters and discarded from downstream analysis. The Hallmark interferon alpha response gene set was procured from the human Molecular Signatures Database (MSigDB), and the AddModuleScore function from Seurat was used to calculate module scores of the gene set for each cell.

Immune response enrichment analysis (IREA) was performed as described.^35^ Briefly, a list of statistically upregulated genes (adjusted p-value < 0.01) in SMNP macrophages compared to alum macrophages was identified using the FindMarkers function in Seurat. This gene list was mapped onto a published scRNA-seq dataset with annotated macrophages treated with 66 different cytokines and PBS ^35^ to characterize how SMNP polarized macrophages. For each comparison, two sets of scores were calculated by summing the normalized expression values of genes in the SMNP gene list in each of the cytokine-treated macrophages and PBS-treated macrophages, and two-sided Wilcoxon rank-sum tests between the two sets of scores were used to assess statistical significance. An FDR correction is applied to all cytokine calculations.

### Statistics

All graphs were prepared in GraphPad Prism Version 10.2.2 and statistical testing was performed using Prism. In the bar graphs, each symbol represents an individual monkey. Datasets were tested for normality and datasets showing lognormal distributions were transformed to linear scale before performing statistical analysis. Statistical comparisons were performed by Student’s *t* test or one-way ANOVA followed by a post hoc test as specified in figure legends. Statistical significance was determined at levels of *p < 0.05, **p < 0.01, and ***p < 0.001.

## Acknowledgements

This work was supported in part by the NIH (awards AI176533 and AI125068 to DJI, award AI048240 to DJI and FV, and award UM1AI144462 to DRB, WRS, and DJI), and the Ragon Institute of MGH, MIT, and Harvard.

## Author Contributions

DJI, FJV, PY, YJZ, LM, and MBM designed the studies. PY, LM, MBM, NL, KKM, and KQ carried out flow cytometry analyses and antibody titer studies. MAR assisted with NHP immunization and carried out PET imaging. PX carried out B cell ELISpot assays. WRS provided immunogens for the studies. MS produced the SMNP adjuvant. LH and RN completed the neutralization assays. SCK carried out analysis of the PET images. LM completed the cell composition analysis. YJZ completed the cell sorting, and sequencing analysis. PY, YJZ, LM, and DJI wrote the manuscript. All authors reviewed and edited the manuscript. JCL, JF, DRB, WRS, FJV and DJI supervised the studies.

## Data availability

Single-cell RNA-seq data, including raw sequencing reads, processed count matrices, and annotated Seurat objects, have been deposited at GEO with the accession number GSE264685 and the reviewer access number kzorcyewbrefril.

