## SUPPLEMENTARY MATERIALS for "Modulation of antigen delivery and lymph node activation in non-human primates by saponin adjuvant SMNP"

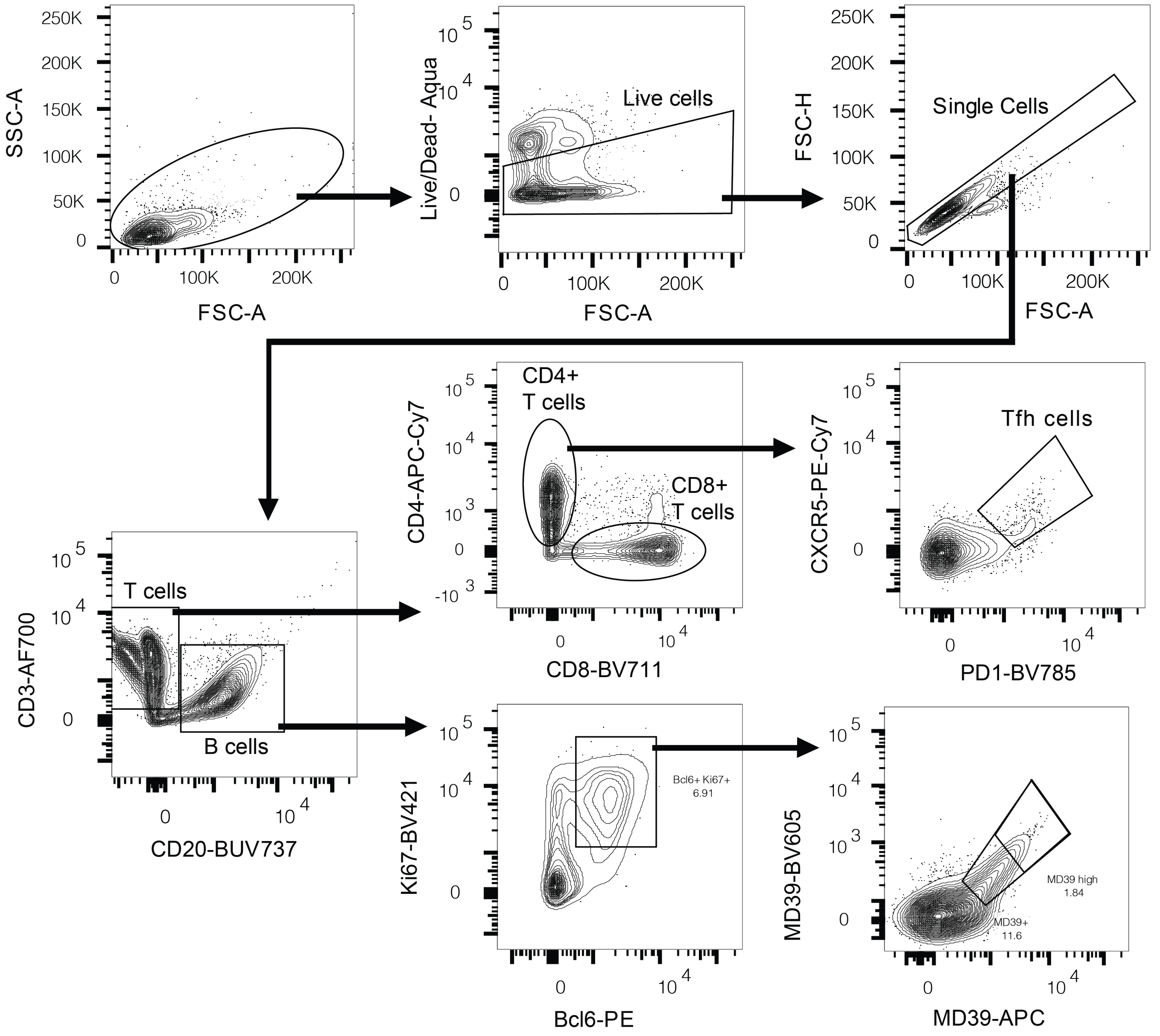


**Figure S1: Flow cytometry gating strategy for analysis of GC Tfh and antigen-specific B cells**. Lymphocytes isolated from FNAs were immunolabeled for the markers indicated. Live cells (Zombie Aqua negative), pre-gated on FSC-A/FSC-H to identify singlets, were gated on CD3 and CD20 markers. B cells (CD3- CD20+) were gated on Ki67 and BCl-6 high expression to identify GC B cells and were then gated twice on MD39-tetramers for antigen-specificity. T cells (CD3+ CD20-) were gated PD1 and CXCR5 high expression to identify Tfh cells.

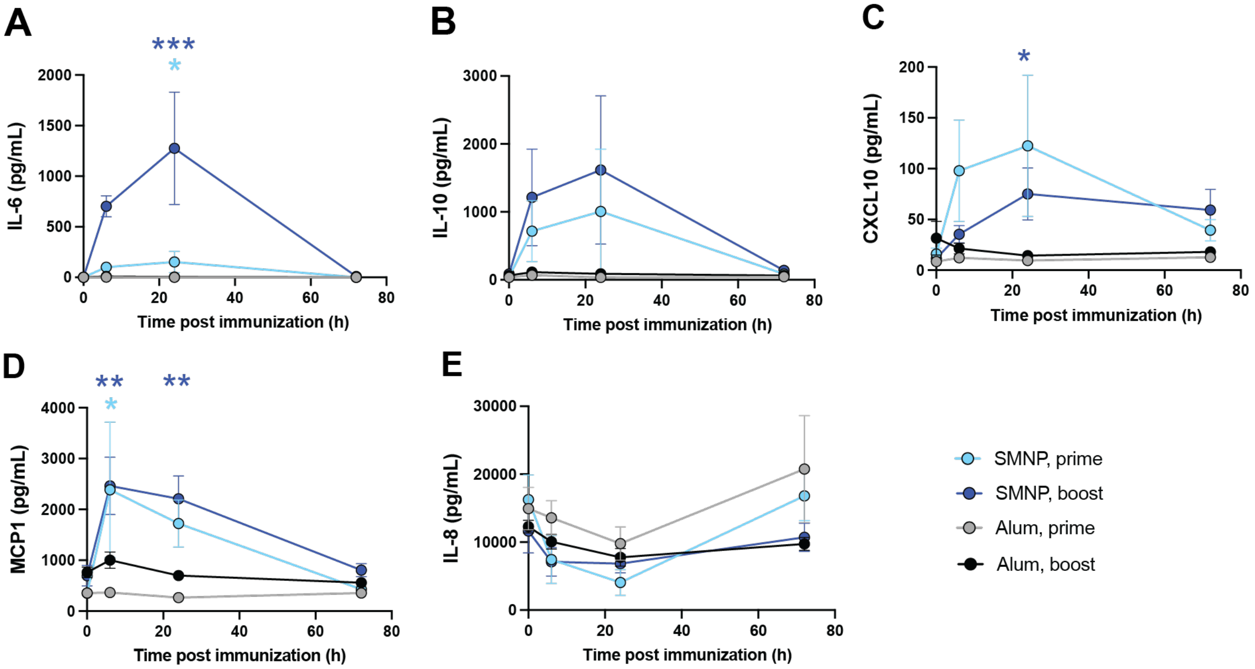


**Figure S2: Cytokine responses to vaccination.** The concentration of cytokines and chemokines in serum was measured at 6, 24, and 72 hours after the prime and first boost immunization. Cytokines analyzed included IL-6 (A), IL-10 (B), CXCL10 (C), MCP1 (D), and IL-8 (E). Statistical analyses were performed using one-way ANOVA, followed by Sidak's post-hoc test. (*P < 0.05, **P < 0.01, ***P < 0.001).

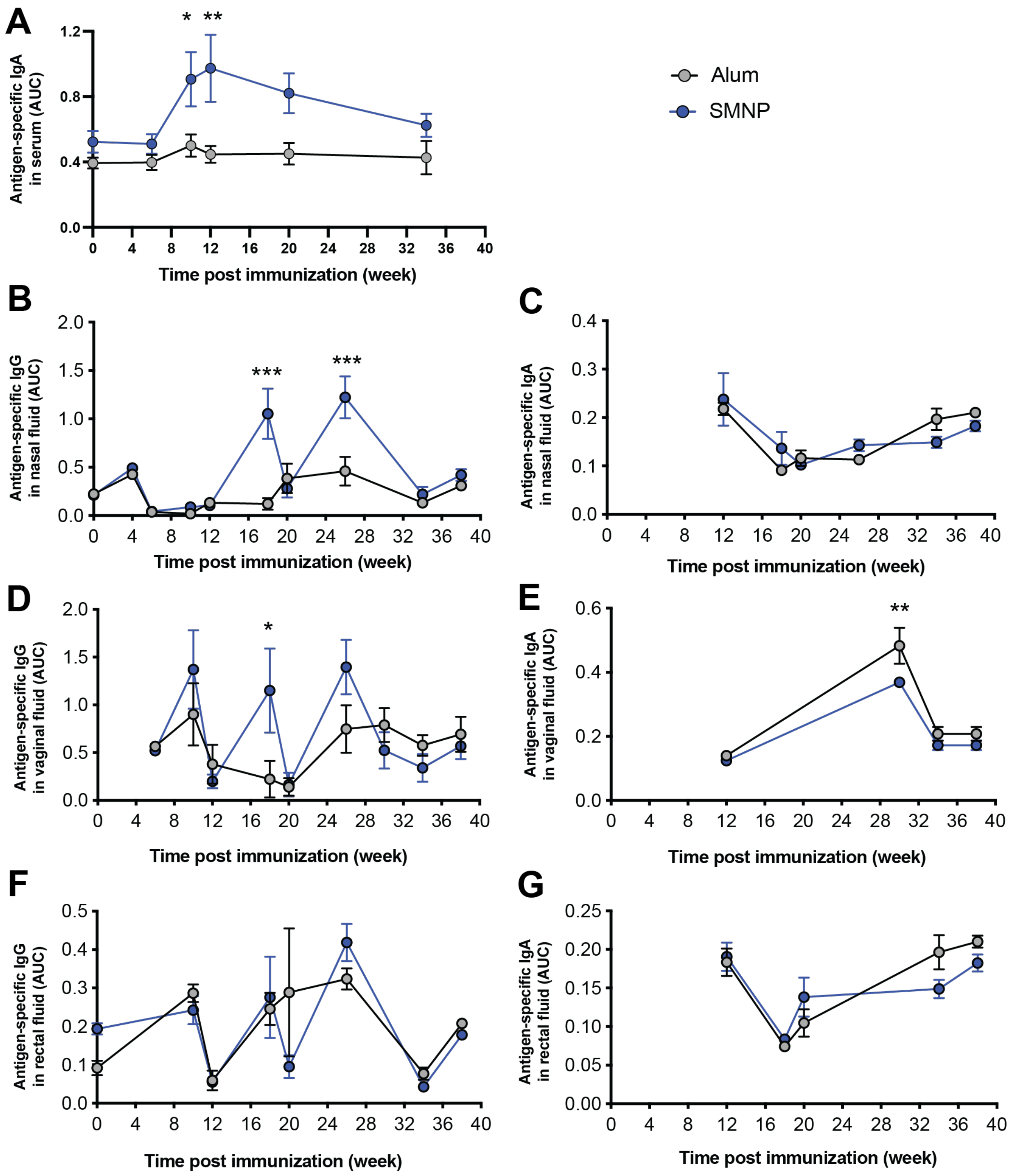


**Figure S3: Systemic and mucosal antibody responses to vaccination.** IgA and IgG were measured in serum and mucosal washes from the (A) nose, (B), (C) vagina and (D) the rectum. Data are presented as mean ± SEM. Statistical comparison between the two groups at each time point was performed by one-way ANOVA followed by Sidak's m post-hoc test was used.

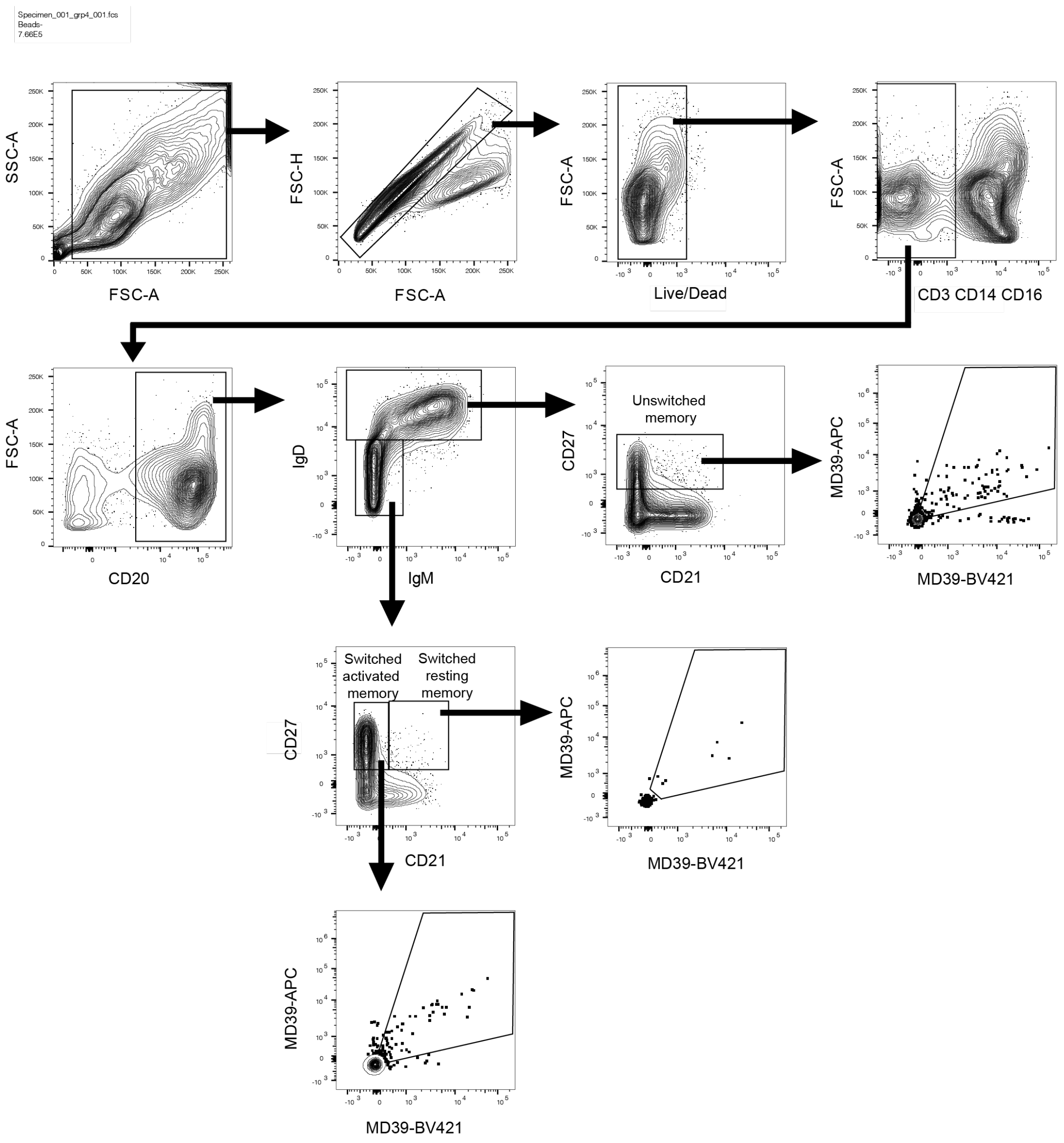


**Figure S4: Flow cytometry gating strategy for analysis of antigen-specific memory B cells.** PBMCs were collected at week 34 and immunolabeled for the markers indicated. Live cells (Zombie Aqua negative), pre-gated on FSC-A/FSC-H to identify singlets, were negatively gated on CD3, CD14, and CD16 markers. Subsequent gating was applied to delineate unswitched, activated switched, and resting switched memory B cells, using markers CD20, IgM, IgD, CD21, and CD27. Lastly, memory B cells were selectively gated twice using MD39-tetramers to determine their antigen specificity.

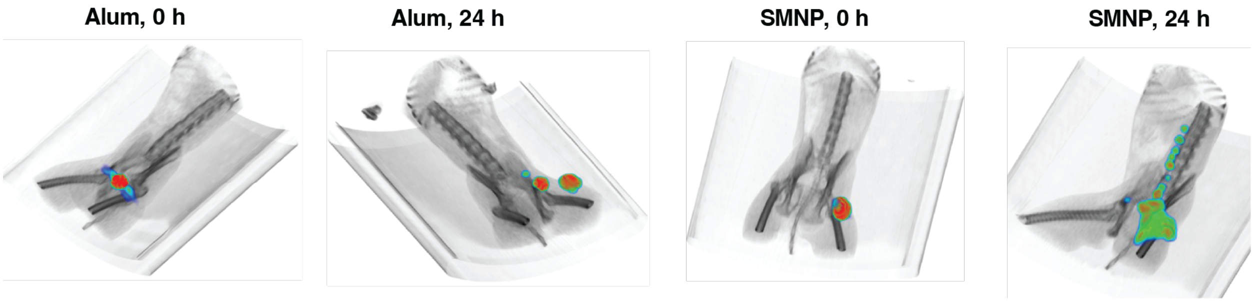


**Figure S5**: Trafficking of SMNP- and alum-adjuvanted MD39 immunogen to draining lymph nodes of macaques. Shown are representative 3D-rendered PET/CT images of the alum-adjuvanted immunization at 0 and 24 h post injection.

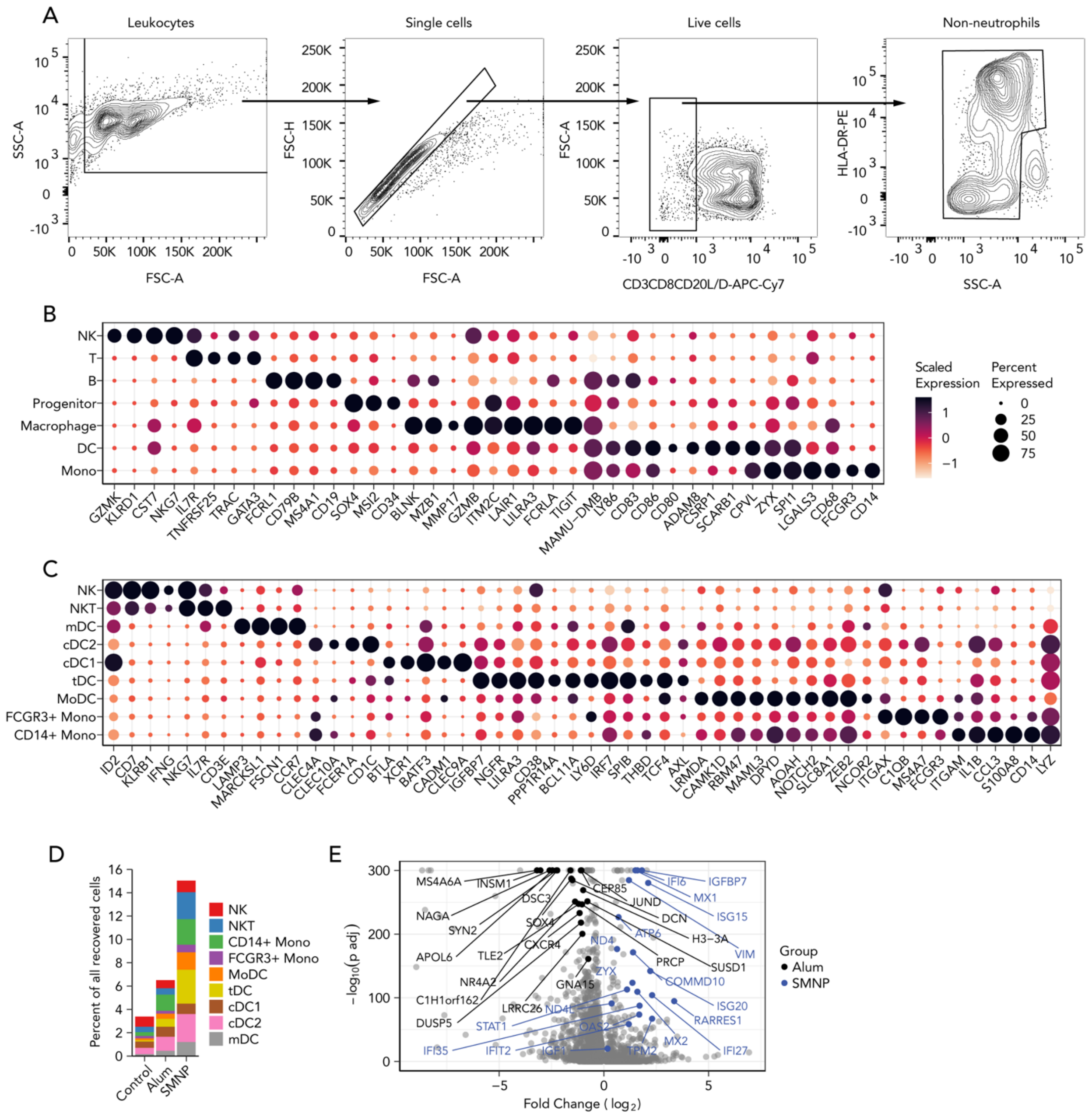

**Figure S6**: **SMNP creates an inflammatory environment in the LN**. (A) Flow cytometry gating strategy used for sorting out CD3$^{low}$, CD8$^{low}$, and CD20$^{low}$ cells for scRNAseq. (B) Dot plot of gene signatures for each cell lineage. (C) Dot plot of gene signatures for the phenotypes of NK cells, DCs, and monocytes. For (B-C), the color of the dots indicates scaled expression levels, and the size of the dots represents the fraction of cells in the cluster that expresses the gene. (D) The percent of recovered cell phenotypes of NK cells, DCs, and monocytes from each group. (E) Volcano plot of differentially expressed genes between SMNP (blue) and alum (black) macrophages.

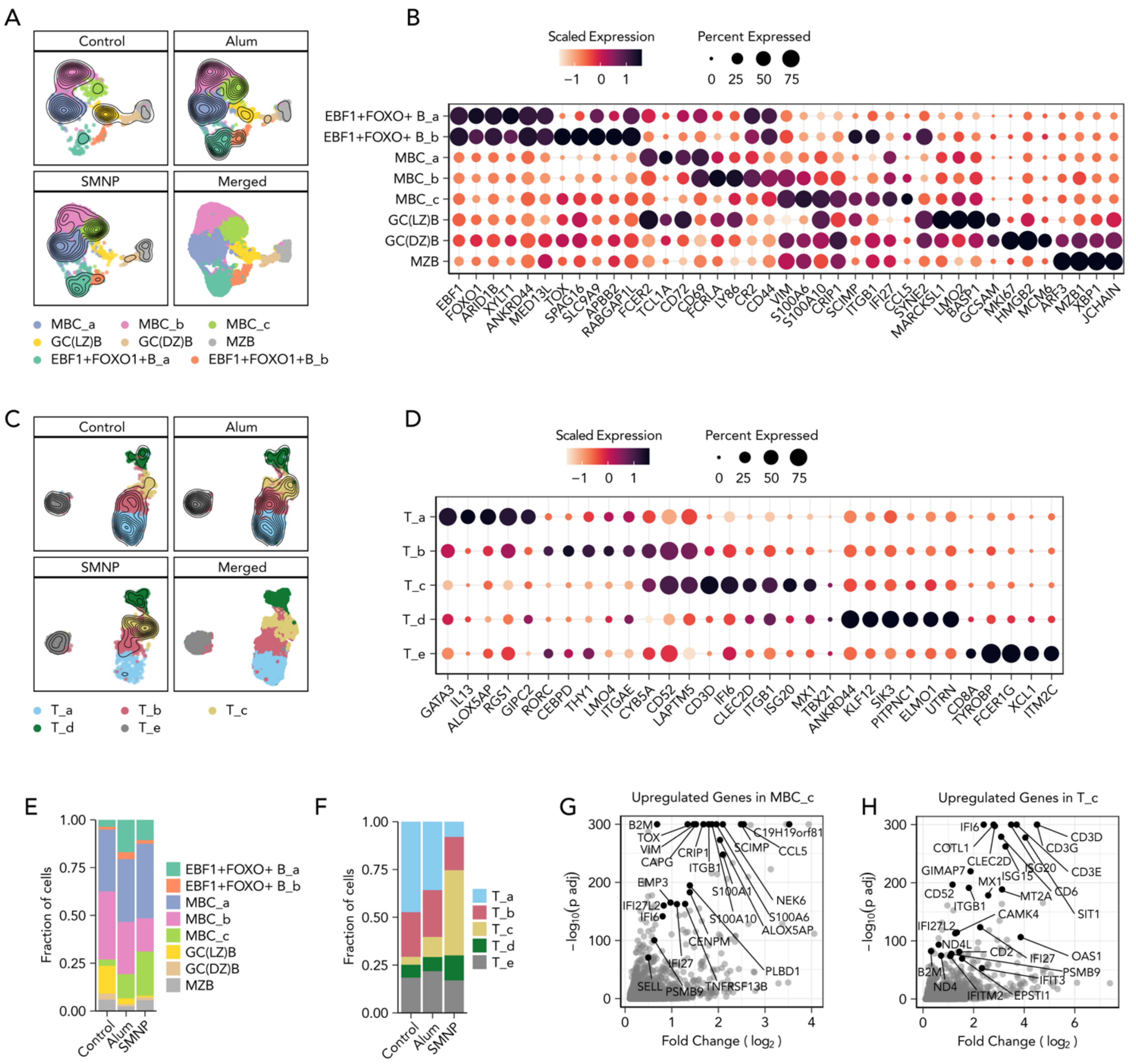


**Figure S7. Transcriptional changes in T and B cell populations in response to alum vs. SMNP immunization**. (A) UMAP of identified B cell phenotypic clusters. (B) Dot plot of gene signatures for each B cell phenotype. (C) UMAP of identified T cell phenotypic clusters. (D) Dot plot of gene signatures for each T cell phenotype. For (A & C), the contour lines represent the density of overlapping cells. For (B & D), the color of the dots indicates scaled expression levels, and the size of the dots represents the fraction of cells in the cluster that expresses the gene. (E) Distribution of identified B cell phenotypic clusters per group. (F) Distribution of identified T cell phenotypic clusters per group. (G) Volcano plot of significantly upregulated genes in MBC_c cluster compared to the rest of B cells. (H) Volcano plot of significantly upregulated genes in T_c cluster compared to the rest of T cells.

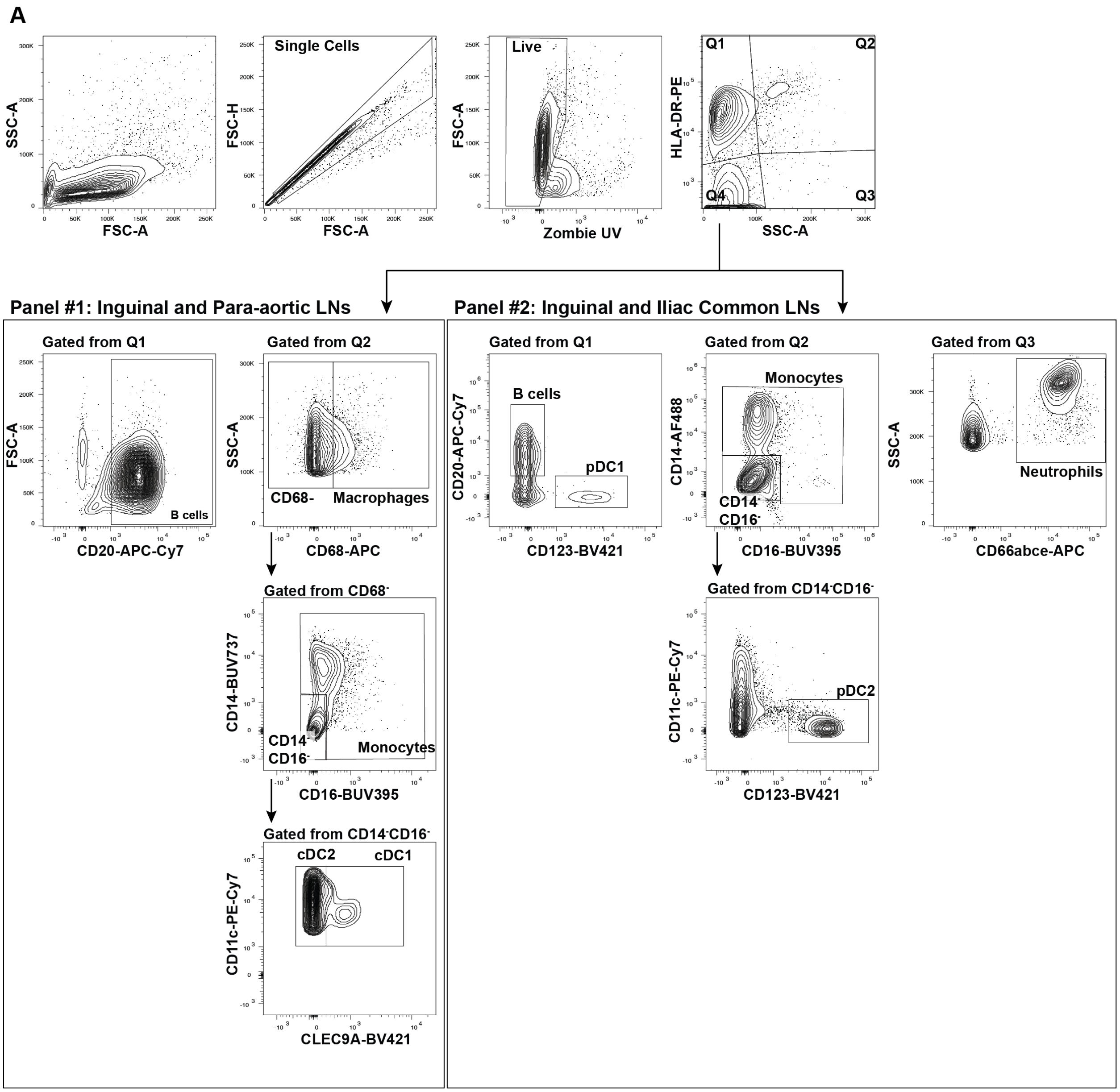


**Figure S8: Flow cytometry gating strategy for analysis of myeloid cells and B cells** in lymph nodes, Live cells (Zombie Aqua negative), pre-gated on FSC-A/FSC-H to identify singlets, were gated on HLA-DR and SSC-A, and then on CD14, CD16, CD20, CD66, CD11c, CD68, and CLEC9A markers to delineate B cells, pDC1 and pDC2 cells, monocytes, neutrophils, macrophages, and cDC1 and cDC2 cells.

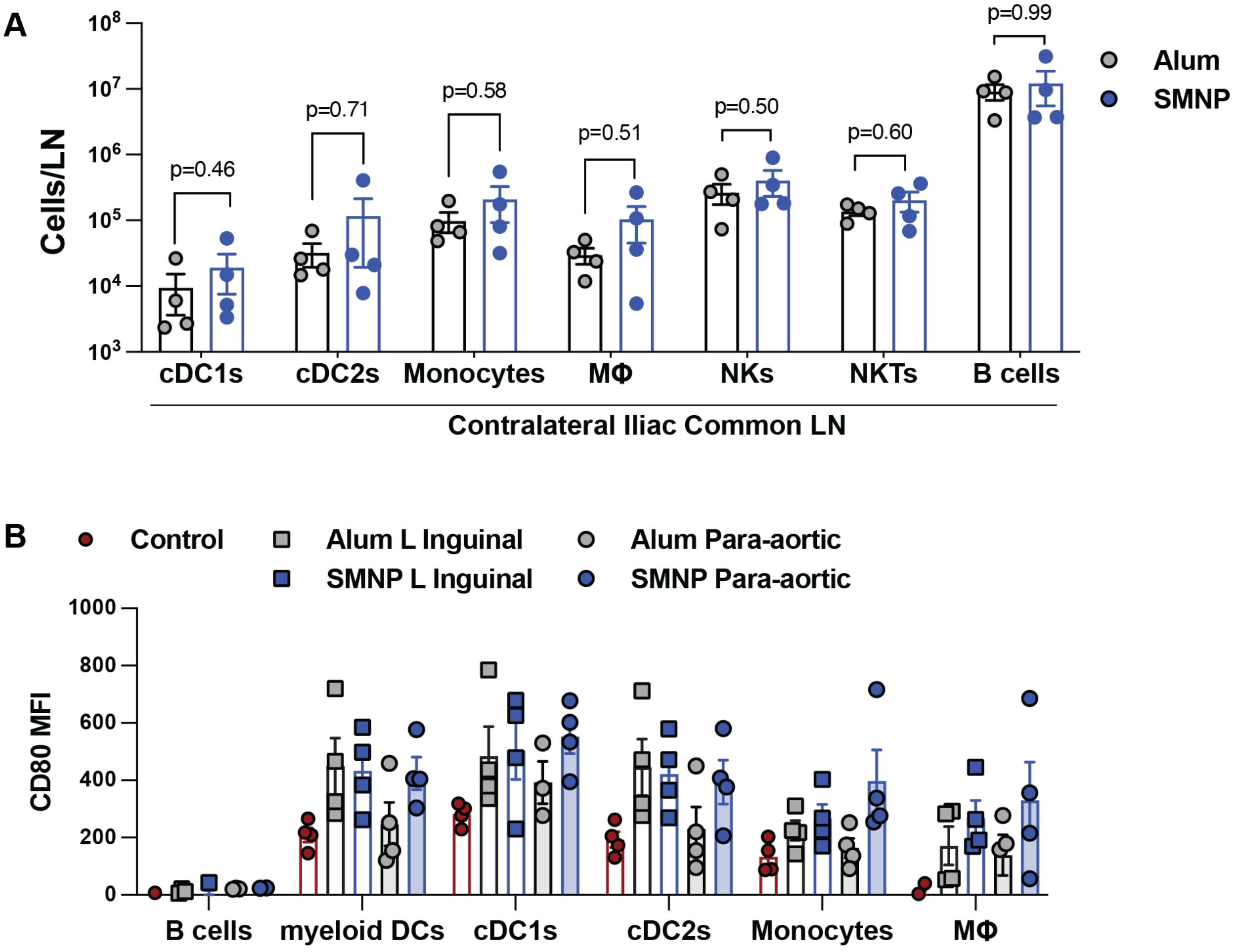


**Figure S9. Innate immune responses to alum vs. SMNP immunization.** (A) Innate immune cell population in control contralateral iliac lymph nodes. (B) CD80 expression on antigen presenting cells in proximal inguinal and distal para-aortic lymph nodes. Statistical analysis was done by Student's *t* test.
